# Prevalence of heterotrophic methylmercury detoxifying bacteria across oceanic regions

**DOI:** 10.1101/2021.08.09.455674

**Authors:** Isabel Sanz-Sáez, Carla Pereira García, Andrea G. Bravo, Laura Trujillo, Martí Pla i Ferriol, Miguel Capilla, Pablo Sánchez, Rosa del Carmen Rodríguez Martín-Doimeadios, Silvia G. Acinas, Olga Sánchez

## Abstract

Microbial reduction of inorganic divalent mercury (Hg^2+^) and methylmercury (MeHg) demethylation is performed by the *mer* operon, specifically by *merA* and *merB* genes respectively, but little is known about the mercury tolerance capacity of marine microorganisms and its prevalence in the global ocean. Here, we explored the distribution of these genes in 290 marine heterotrophic bacteria (*Alteromonas* and *Marinobacter* spp.) isolated from different oceanographic regions and depths, and assessed their tolerance to diverse concentrations of Hg^2+^ and MeHg. About 25% of the isolates presented *merA* and only 8.9% presented both *merAB* genes, including the strain ISS312 that exhibited the highest tolerance capacity and a degradation efficiency of 98.2% in 24 h. Fragment recruitment analyses of ISS312 genome against microbial metagenomes indicated an extensive distribution across the global bathypelagic ocean. Our findings highlighted that mercury resistance genes are widely distributed in a non-highly polluted environment such as the pelagic marine environment, and that degradation of the neurotoxic MeHg can be performed through the ocean water column by some heterotrophic bacteria at high efficiency with important implications in the biogeochemical cycle of mercury and potentially for the environment and human health.

**Teaser:** Active mercury resistance genes detected in marine cultured bacteria are widely distributed in the ocean including the bathypelagic zone.

## 1. Introduction

Mercury (Hg) is one of the most toxic, widespread and worrisome contaminants (*1*, *2*), which is emitted to the atmosphere by natural sources, such as volcanoes and rock weathering, but also by anthropogenic activities. Indeed, it has been estimated that anthropogenic Hg emissions have enriched present-day atmosphere by a factor of 2.6 relative to 1840 levels, and by a factor of 7.5 relative to natural levels (*3*). The rising Hg levels since the industrial era, estimated as an increase of 450 % of Hg in the atmosphere (*4*), makes the study of its biogeochemical cycle a major concern to the scientific community and also to all the governments around the world. The Minamata convention, held in 2013 and entering into force in August 2017 (*5*). has been created to regulate the use of Hg and its releases to the environment in order to reduce its current levels.

Emitted elemental (Hg^0^) and inorganic divalent Hg (Hg^2+^) can be deposited to land and oceans by wet and dry depositions (*6*, *7*). Hg^2+^ in the ocean can be then volatilized back again to the atmosphere as Hg^0^ (*8*), or can be methylated (*9–13*) forming methylmercury (MeHg), which bioaccumulates and biomagnifies in aquatic food webs (*4*, *14*, *15*). As a consequence, humans are exposed to this neurotoxicant mainly through fish and seafood consumption (*14*, *16*, *17*). MeHg levels in the oceans vary with depth, and usually, measures are being reported low in open ocean surface waters, maximal in intermediate layers, especially in regions of low-oxygen and near or below the thermoclines (up to 1000 m depth), and low and relatively constant in deeper waters (>1000 m depth) (*14*, *18*, *19*). While Hg^2+^ methylation has been reported to occur in oxic and sub-oxic layers of the water column (*9*, *18*, *20–22*) mainly associated with the microbial remineralization of sinking particulate organic matter (*18*, *19*, *23*), much less is known about MeHg demethylation and Hg^2+^ reduction processes in the ocean water column. Although MeHg demethylation and Hg^2+^ reduction processes can be photochemically mediated (*24–26*), light penetration in the ocean water column is limited to 200 m (*27*), thereby biological MeHg degradation and Hg reduction processes likely govern in the ocean water column.

Biological MeHg demethylation and Hg^2+^ reduction detoxification processes are mediated by the *mer* operon (*28*, *29*). While the operon can be composed by different sets of genes (*28*, *29*), the operon key genes are *merA* and *merB*. The first one codifies a mercuric reductase and is responsible for the transformation of Hg^2+^ to the less harmful and volatile Hg^0^ (*28*). The *merB* gene encodes an organomercurial lyase enzyme that confers resistance to the organic MeHg form. It is the responsible for its demethylation releasing Hg^2+^ that will be then reduced to Hg^0^ by *merA* gene (*28*). These machineries have been found in numerous microorganisms including aerobic and anaerobic species, although demethylation appears to be predominantly accomplished by aerobic organisms (*29*, *30*). To date, very few studies have reported the presence of *mer* genes in oceanic waters, with the exception of some studies in the North Pacific and Arctic Oceans (*31*) and a recent analysis in the global bathypelagic ocean (Bravo & Sanz et al. unpublished). Taking into account that different concentrations of MeHg can be found through the ocean water column (*21*, *32–34*), it would be however plausible to find microorganisms with the Hg detoxification capacity. In this study we took advantage of the MARINHET (*35*) culture collection, that includes a total of 2003 marine bacterial strains from a wide variety of oceanographic regions and depths, to perform a functional screening of the *merA* and *merB* genes in 290 marine heterotrophic bacteria with the aim to detect marine bacteria codifying *mer* genes. Moreover, we assessed their tolerance to different concentrations of inorganic (Hg^2+^) and organic monomethylmercury (MeHg).

Most of the studies focusing on the isolation of marine Hg resistant bacteria have targeted coastal seawaters (*36–38*), sediments (*37*, *39*), mangroves and estuaries (*40–42*) hydrothermal vents (*43*), and specially highly contaminated sites (*44*, *45*) and have overlooked the open ocean including the bathypelagic ocean. Identification of Hg resistant bacteria in contrasting aquatic ecosystems and the assessment of their tolerance to different concentrations of MeHg provides new opportunities to explore the ubiquity, prevalence of marine cultured bacteria with detoxification capacity in the open ocean (i.e. non-contaminated sites), and also, it appears to be an interesting starting point for bioremediation strategies.

## 2. Results and Discussion

### 2.1 Presence of merA and merB genes among Alteromonas and Marinobacter strains

Hg resistance genes have been found in multiple Gram-negative and Gram-positive bacteria, and isolates containing the *mer* operon have been retrieved from many environments (*46–52*), including different marine ecosystems (*36*, *43*, *53*, *54*). However, most of the studies have found them in highly contaminated environments and their presence in pelagic open ocean environments have been largely overlooked. We used the MARINHET bacterial culture collection (*35*), which included strains from different depths, such as the surface, the deep chlorophyll maximum and the bathypelagic zone, as well as from diverse oceanographic regions, to explore the presence of the *mer* operon in non-contaminated sites. We detected a total of 20 different bacterial taxa in the IMG/JGI database that matched at the genera level with the taxonomic assignation of the MARINHET culture collection isolates, and therefore, with potential for carrying *merA* and *merB* genes (**Supplementary Table S1**). The comparative analyses between the 16S rRNA gene sequences of these 20 genera containing the targeted *merA* and *merB* genes and the partial 16S rRNA sequences of our isolates revealed a total of 352 strains that were, at least, 99 % identical to one of the putative candidates genera. These comprised 7 genera (*Alteromonas*, *Marinobacter*, *Idiomarina*, *Pseudomonas*, *Micrococcus*, *Zunongwangia* and *Bacillus*) (**Supplementary Table S2**). From these, we selected a total of 244 strains affiliating to *Alteromonas sp*. and 46 strains to *Marinobacter sp*. (**Supplementary Table S3**). These two genera were chosen for *merA* and *merB* functional screening for several reasons: (i) they are among the most common culturable heterotrophic bacteria living in open marine waters all around the world, as they have been isolated from a wide variety of marine environments (*55–61*), and in the case of *Alteromonas* it is one of the most ubiquitous cultured taxa in the ocean (*35*), (ii) it has been already described that species of those genera harbor in their genomes the *mer* operon (*59*, *62–65*) and finally (iii) they were found highly abundant in our MARINHET culture collection.

The functional screening of the *merA* and *merB* genes from the 244 *Alteromonas* and 46 *Marinobacter* strains revealed that 13.5 % (32 out of 244) and 89.1 % (41 out of 46) of the strains presented only *merA*, respectively, while only 1.6 % (4 out of 244), and 47.8 % (22 out of 46) presented both *merA* and *merB* genes (*merAB*) (**Table 1**). These results showed that *Marinobacter* displayed a higher proportion of *merAB* genes than *Alteromonas*. In addition, our results showed that the *merB* gene was found in lower proportion than *merA* among both *Alteromonas* and *Marinobacter* strains, being these results consistent with the known fact that the *mer* operon does not always codify for the organomercurial lyase necessary for the detoxification of organic Hg compounds (*28*, *29*). In addition, studies targeting the abundance of *mer* genes in the environment reported that *merA* is found to be widely distributed among marine bacteria (*29*, *31*, *66*), while *merB* only has an ecological significance in determined systems where MeHg is present at high concentrations (*67*). Here, we demonstrate that both *merA* and *merB* might be present even in pristine environments such as the open ocean.

**Table 1.** Summary of the PCR screening results for *merA* and *merAB* in *Alteromonas* and *Marinobacter* strains. Photic includes surface and deep chlorophyll maximum (DCM) isolates, while aphotic includes bathypelagic isolates.

| Genus | Layer | N° of tested strains | Positives PCR for |  | Total strains with *merA* and/or *merAB* |
| --- | --- | --- | --- | --- | --- |
|  |  |  | *merA* | *merAB* |  |
| *Alteromonas* | Photic | 127 | 13 (10.2%) | 1 (0.8%) | 33<br>(13.5%) |
|  | **Aphotic** | 117 | 19 (16.2%) | 3 (2.6%) |  |
| *Marinobacter* | Photic | 33 | 30 (90.9%) | 20 (60.6%) | 41<br>(89.1%) |
|  | **Aphotic** | 13 | 10 (76.9%) | 2 (15.4%) |  |

### 2.2 Biogeographic distribution of isolated strains codifying mer genes

The strains screened for *merAB* genes were isolated from different oceanic regions such as the North Western Mediterranean Sea (89), South (101) and North Atlantic (42), Indian (44), Arctic (7) and Southern (7) Oceans and included isolates from photic (160) and aphotic (130) layers (**Table 2**). We obtained positive strains (*merA* and/or *merAB*) from the different depths and oceanographic regions, except for the Arctic Ocean (**Figure 1 and Supplementary Table S4**). For *Alteromonas* sp., station (ST) 32 from the bathypelagic in the South Atlantic Ocean and ST151 from surface of the North Atlantic Ocean were the ones with a larger number of positive strains (**Figure 1**), while ST76 and ST67 from surface South Atlantic Ocean followed by ST8 of the bathypelagic North Western Mediterranean Sea were the ones which presented more positive *Marinobacter* strains (**Figure 1**). No significative differences were found between oceans (ANOVA, P-value > 0.05) but in general, we found a higher proportion of positive strains coming from waters of the Southern Ocean (71 % despite the lower number of strains tested) and the South Atlantic Ocean (48 %), followed by those retrieved from the North Atlantic (17 %) and North Western Mediterranean Sea (13 %). However, total Hg concentrations have been recorded to be higher in the Mediterranean and the North Atlantic Ocean compared to the South Atlantic and Southern Oceans, where concentrations were lower (*34*). Nevertheless, some studies have highlighted the important but also variable levels of MeHg found in Southern and other polar waters compared to open ocean (*32*, *68*), and isolation of Hg resistance bacteria from those polar waters has been previously reported (*52*). It must be noticed that our results only represent a minor fraction of all potential isolates that may harbor *merAB* genes, since our primers only targeted *Alteromonas* and *Marinobacter* genera, and only specific *merA* gene variants. Thus, correlation between levels of Hg and positive strains cannot be properly assessed.

**Figure 1.**
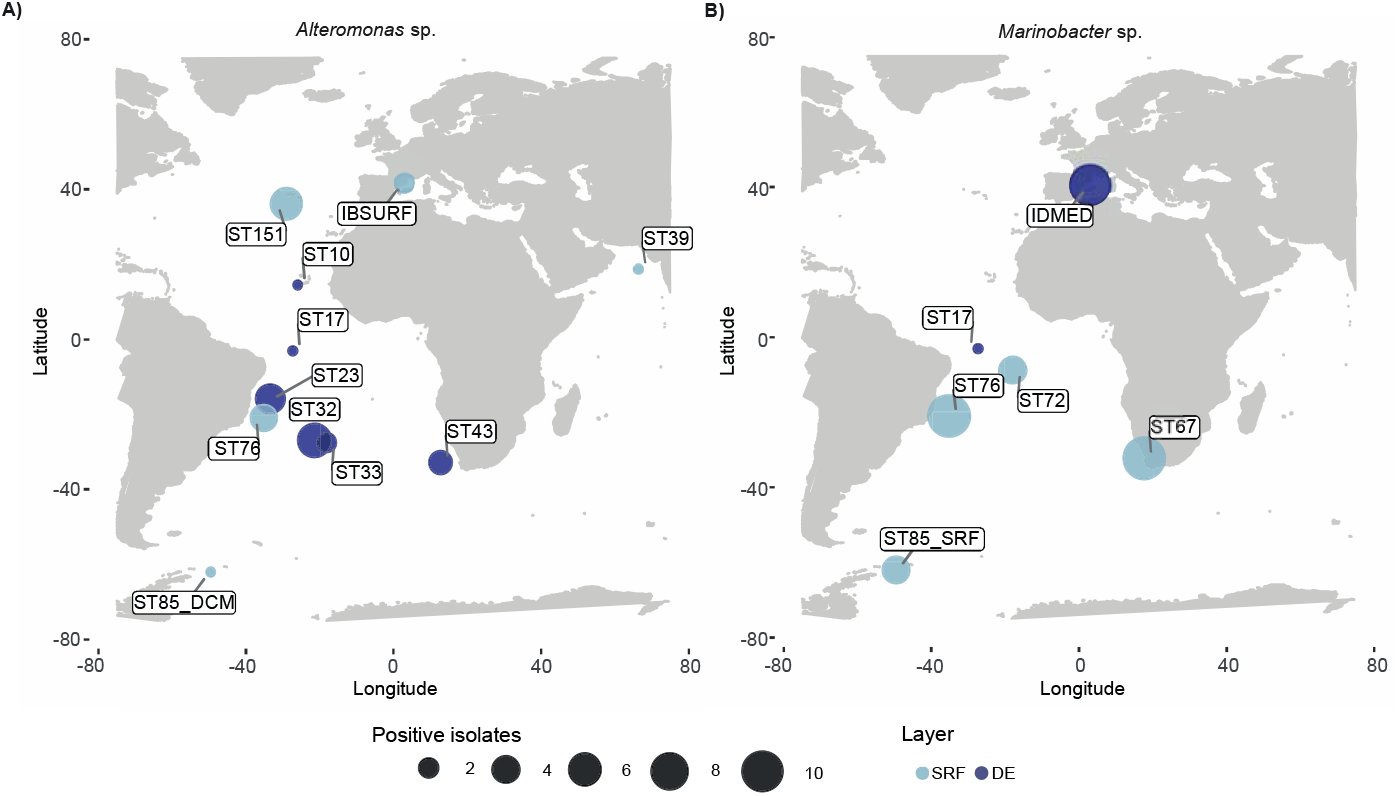
Distribution of the positive strains after functional screening for *merA* and/or *merAB* genes. **(A)** *Alteromonas* sp. strains **(B)** *Marinobacter* sp. strains. Size of the dots indicates how many strains per station including *merA* and/or *merAB* were identified. Color of the dots indicated the layer (photic or bathypelagic) where the strains were retrieved.

**Table 2.** Characteristics of the marine seawater samples from five different cruises used for the isolation of bacteria screened for Hg resistance genes. ATP09: Arctic Tipping Points cruise in 2009; BBMO: Blanes Bay Microbial Observatory.

| **Cruise** | **Station** | **Oceanic location** | **Latitude** | **Longitude** | **Depth (m)** | **N° of screened isolates** | **N° of *merA* positives** | **N° of *merAB* positives** |
| --- | --- | --- | --- | --- | --- | --- | --- | --- |
| *Tara*<br>Oceans | ST 39 | Indian Ocean | 18° 35.2' N | 66° 28.22' E | 25 | 44 | 0 | 1 |
|  | ST 67 | South Atlantic | 32° 17.31' S | 17° 12.22' E | 5 | 20 | 11 | 6 |
|  | ST 72 | South Atlantic | 8° 46.44' S | 17° 54.36' W | 5 | 15 | 4 | 2 |
|  | ST 76 | South Atlantic | 20° 56.7' S | 35° 10.49' W | 5 | 31 | 15 | 8 |
|  | ST 85 | Southern Ocean | 62° 2.19' S | 49° 31.44' W | 5.9 | 1 | 1 | 0 |
|  | ST 85 | Southern Ocean | 62° 2.19' S | 49° 31.44' W | 87.4 | 6 | 4 | 4 |
|  | ST 151 | North Atlantic | 36° 10.17' N | 29° 1.23' W | 5 | 31 | 6 | 0 |
| ATP09 | AR_1 | Arctic Ocean | 78° 20.00' N | 15° 00.00' E | 2 | 3 | 0 | 0 |
|  | AR_2 | Arctic Ocean | 76° 28.65' N | 28° 00.62' E | 25 | 4 | 0 | 0 |
| Malaspina | ST 10 | North Atlantic | 21° 33.36' N | 23° 26' W | 4002 | 11 | 1 | 0 |
|  | ST 17 | South Atlantic | 3° 1.48' S | 27° 19.48' W | 4002 | 3 | 2 | 0 |
|  | ST 23 | South Atlantic | 15° 49.48' S | 33° 24.36' W | 4003 | 13 | 5 | 0 |
|  | ST 32 | South Atlantic | 26° 56.8' S | 21° 24' W | 3200 | 14 | 7 | 0 |
|  | ST 33 | South Atlantic | 27° 33.2' S | 18° 5.4' W | 3904 | 2 | 2 | 2 |
|  | ST 43 | South Atlantic | 32° 48.8' S | 12° 46.2' E | 4000 | 3 | 3 | 2 |
| MIFASOL | ST 8 | NW Mediterranean | 40° 38.41' N | 2° 50' E | 2000 | 84 | 9 | 2 |
| BBMO | IBSURF | NW Mediterranean | 41° 40' N | 2° 48' E | 5 | 5 | 2 | 0 |

Regarding the presence of the *merA and merAB* genes across the ocean layers, isolates came from both depths and 27 % and 23 % of the total surface and the deep-ocean isolates screened, respectively, gave positive results. Although, no significative differences were found between depths (ANOVA, P-value >0.05) our findings unveiled the pattern of vertical distribution of these *merAB* genes distributed along different water depths including bathypelagic waters.

### 2.3 High variability of mercury tolerance within marine bacteria

It is unknow whether mercury tolerance is a conservative trait within marine bacterial strains of the same genera. MIC experiments were addressed in a selection of the 74 isolates that presented *merA* and/or *merAB* genes. This selection was based on a clustering of the 16S rRNA gene of the isolates at 99 % sequence similarity to define operational taxonomic units (OTUs). The clustering grouped the isolates into 8 OTUs, and representatives of different OTUs were randomly selected for MIC experiments (**Supplementary Table S5**). First, we tested the tolerance for inorganic mercury (HgCl_2_) and 42 isolates (19 *Alteromonas* and 23 *Marinobacter*) displayed different levels of tolerance. MIC values ranged generally from 10 to 50 μM. Around 50 % of the *Alteromonas* and *Marinobacter* strains tested presented a MIC of 20 μM and one of the isolates stood out as it presented a tolerance to HgCl2 up to 70 μM (**Supplementary Table S6**). Different tolerances were found within the same phylogenetic cluster based on 16S rRNA genes (**Figure 2**) but also within strains belonging to the same OTUs (99% sequence similarity, **Figure 2**). For example, within the cluster of *Alteromonas mediterranea* some strains presented a MIC of 20 μM, while the isolate that presented the highest tolerance (70 μM, ISS312) also belonged to the same *Alteromonas* species (**Figure 2**). The same occurred among *Marinobacter* isolates, where members of the *Marinobacter hydrocarbonoclasticus* cluster presented MIC values ranging from 10 to 50 μM (**Supplementary Table S6 and Figure 2**).

**Figure 2.**
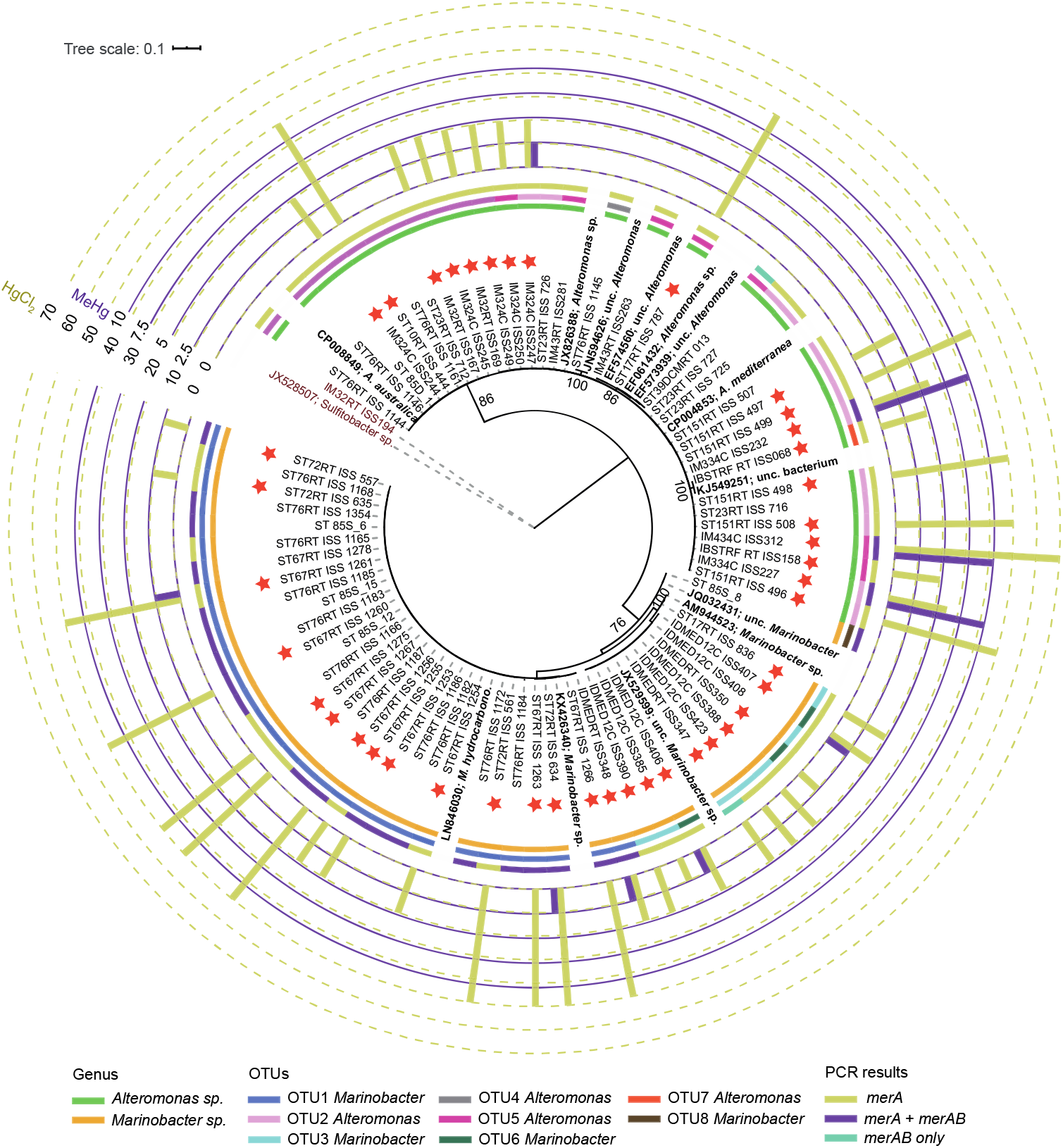
Phylogeny of the 16S rRNA gene of *Alteromonas* and *Marinobacter* positive strains for *merA* and/or *merAB* genes screening. First inner colored strip indicates genus of the strain. Second colored strip indicates to which OTU (99% sequence similarity in their 16S rRNA gene) each of the strains belong. Third colored strip indicates presence or absence of genes based on PCR results. Red stars indicate those isolates where MIC experiments where performed. Bars indicate results from the MIC experiments: yellow, HgCl2; purple, MeHg. Tolerance values are in μM. JX52807, *Sufitobacter* and IM32RT_ISS194 are outgroups of the tree. The numbers in the nodes represent bootstrap percentages > 75%. *A. australica*, *Alteromonas australica*; *A. mediterranea*, *Alteromonas mediterranea*; unc., uncultured; *M. hydrocarbon*., *Marinobacter hydrocarbonoclasticus*.

The amino acids *merA* phylogeny, which theoretically could include the different gene sequences variants covered by the primers designed, unveiled that all the *Alteromonas merA* sequences grouped into two sister clades with the reference *Alteromonas mediterranea merA* sequence **(Figure 3)** and MIC variability was more clearly observed here as sequences clustering together presented MIC values ranging from 20 μM to 70 μM. For *Marinobacter*, the *merA* genes grouped into three different clusters showing genetic heterogeneity among their *merA* genes copies (**Figure 3**). Those sequences of the *Marinobacter* sp. Arc7-DN-1 cluster presented a MIC that varied between 20 μM and 50 μM, while those of the *merA* sequences affiliating to *Marinobacter salarius* and *Marinobacter hydrocarbonoclasticus* ranged from 10 μM to 50 μM **(Figure 3)**.

**Figure 3.**
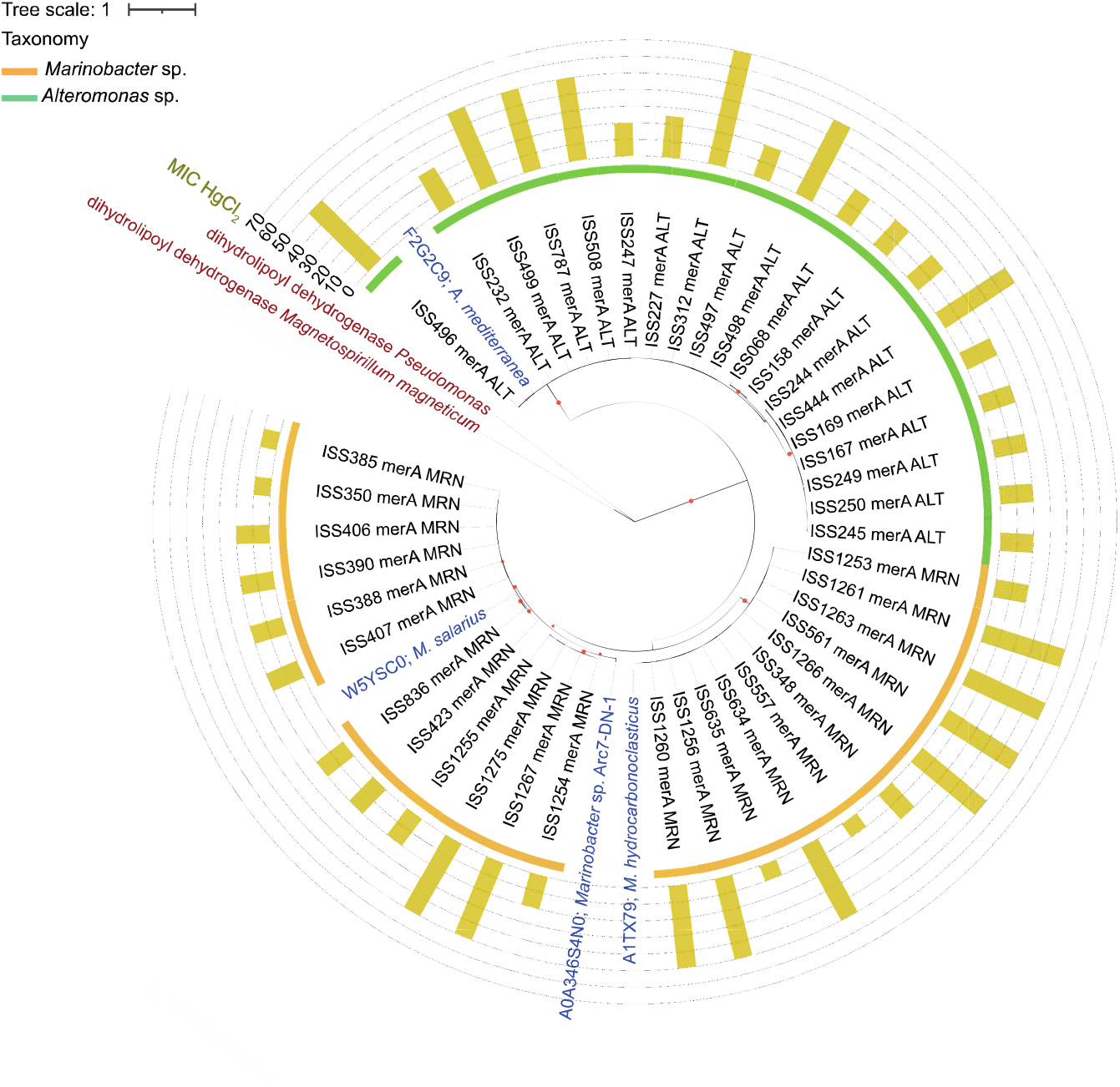
Phylogeny of the *merA* amino acid sequences from *Alteromonas* and *Marinobacter* positive strains. Including only those sequences codified in those strains submitted to Minimum Inhibitory Concentration (MIC) assays. MIC results for the tested strains against HgCl_2_ are indicated by bars. Sequences of dihydrolipoyl dehydrogenase act as outgroups of the tree. Color strip indicates taxonomy of the sequences: *Alteromonas* sp., green; *Marinobacter* sp., orange. Reference sequences are indicated in bold blue: *A. mediterranea*, *Alteromonas mediterranea*; *M. hydro*., *Marinobacter hydrocarbonoclasticus*; *M. salarius*, *Marinobacter salarius*. Bootstrap values >75% are indicated by red circles in the tree nodes.

On the other hand, in order to test the tolerance to MeHg amended in form of CH_3_HgCl, three strains affiliating to *Alteromonas mediterranea* that codified for the *merB* gene and presented a MIC for HgCl_2_ above 20 μM were selected. Remarkably, these strains presented a high tolerance to MeHg, growing at concentrations up to 10 μM (**Figure 3**) and the *merAB* aminoacids sequences clustered all together, as expected, with *Alteromonas mediterranea* reference sequence **(Supplementary Figure S1)**. Among the *Marinobacter* sp. harboring *merAB* genes, the two strains having a MIC for HgCl_2_ above 20 μM did not show a substantial growth above 2.5 μM of MeHg (**Figure 2**).

The MIC values heterogeneity within strains belonging to the same OTU or phylogenetic cluster, suggested that the level of Hg resistance was isolate specific, and that maybe we retrieved different ecotypes within a same species with different tolerances to Hg. In addition, the operon *mer* can be either codified in the chromosome (*69*) or in plasmids (*28*, *70*), and usually, *mer* genes are components of transposons (*71*), and integrons (*72*, *73*). Thus, different copies of the *mer* genes including maybe different sequence variants (*29*, *74*), could be found within a same strain providing to the strains different tolerances to mercury compounds. This specificity is further supported by the fact that no correlation was observed between the ability to reduce Hg and the taxonomic groups observed in the phylogenetic trees, although negative results should be interpreted with caution. Similar results were obtained in previous studies indicating the isolate-specific resistance to Hg (*44*, *75*). Despite these differences between strains, the tolerances found for inorganic Hg were similar to those found in other studies where *Alteromonas* (*40*, *65*, *69*, *76*) and *Marinobacter* (*43*) genera were also isolated from different marine ecosystems such as hydrothermal vents, estuaries or contaminated sediments. However, to the best of our knowledge, this is the first study addressing the tolerance of *Marinobacter spp*. and *Alteromonas mediterranea* isolated from the ocean to MeHg. Hence, we found out that a strain affiliating to *Alteromonas mediterranea* (ISS312) presented a MIC to inorganic Hg higher than other strains already published, up to 70 μM, but we also determined that it was able to grow in the presence of MeHg, presenting a MIC up to 10 μM. It is noteworthy that the tolerant strains to HgCl_2_ and/or to MeHg were resistant to much higher concentrations of Hg than those reported in different oceans, which usually range from <0.1 to 10 pM (*34*, *77*).

### 2.4 Description of the highly tolerant Alteromonas sp. strain ISS312

Strain ISS312, isolated from South Atlantic bathypelagic waters at 4000 m, affiliated to the species *Alteromonas mediterranea*. It displayed the highest tolerance to both HgCl_2_ (70 μM) and MeHg (10 μM) and it could be a good candidate for future bioremediation studies in highly contaminated areas with both organic and inorganic mercuric compounds. Consequently, the growth rates of this isolate at different concentrations of MeHg were assessed. Tested concentrations were selected based on MIC results and included: a control without MeHg (0 μM), 1 μM, 2.5 μM and 5 μM MeHg. Growth curves at 0 μM and 1 μM were very similar, as well as between 2.5 μM and 5 μM (**Figure 4A**). We observed that the major difference between growth curves was the length of the lag phase, where bacteria adapt themselves to the growth conditions. Cultures showed a longer lag phase in the highest concentration of MeHg, around 13 h, compared to the control, which started to grow immediately after inoculation (**Figure 4A**). This phenomenon seems to be a common trait for Hg resistant strains in the presence of the toxic compound, as this behavior has been repeatedly observed in different species of *Pseudomonas* sp., *Alcaligenes* sp. or *Bacillus* sp. (*37*, *53*, *78*). Lag phase length declined as long as the concentration of MeHg decreased, being 6 h at 2.5 μM and 2 h at 1 μM. However, once the cultures started to grow, their growth rates (*μ_maX_*) were very similar independently of their initial MeHg concentrations, ranging from 0.10 h^-1^ in the control to 0.09 h^-1^ at 5 μM. Stationary phase was reached in all concentrations at 80 h, even though at this time cultures at higher concentrations of MeHg seemed to be only entering to the plateau (**Figure 4A**). In addition, their carrying capacity (k), i.e. the maximum population size of a species, was between 1.6 and 1.9 based in O.D. measures, revealing very similar values between tested concentrations, an observation also recurrently reported (*37*, *79*). TEM observations of the ISS312 cultures growing at 0 μM and 5 μM of MeHg also showed similar morphology and ultrastructure of the cells **(Figure 4A)**.

**Figure 4.**
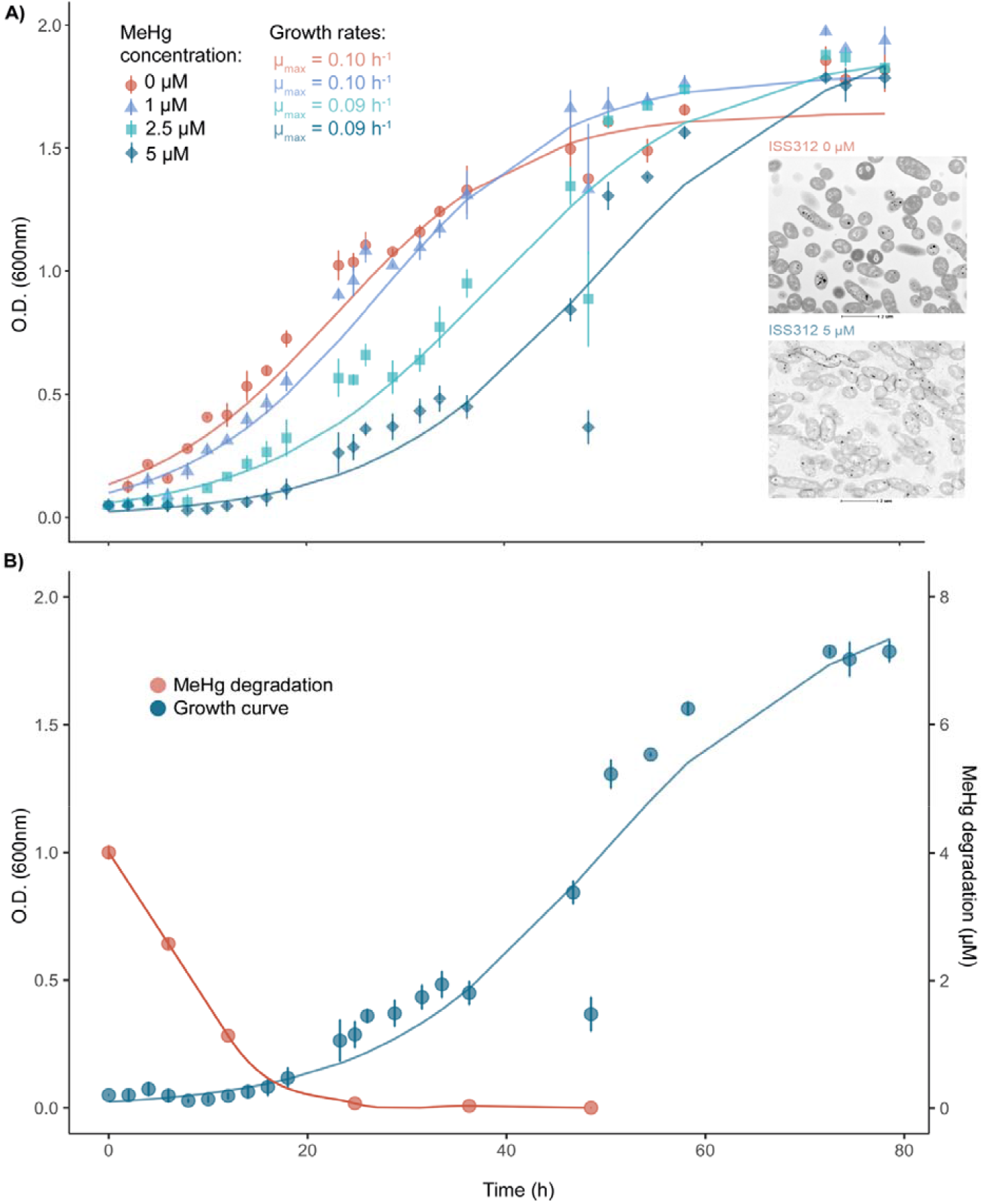
Growth effect of MeHg in strain ISS312. **(A)** Growth kinetics of *Alteromonas mediterranea* strain ISS312 in Zobell broth containing MeHg (control (0 μM), 1 μM, 2.5 μM and 5 μM). *μ_max_* indicates the maximum growth rate for each MeHg concentration. TEM images of the strain growing at 0 μM and at 5 μM are shown in the right side of the plot. **(B)** MeHg removal by strain ISS312 during the growth curve experiment at 5 μM. Mean and standard deviation from three replicates samples are shown in both graphs.

In addition, samples for the measurement of MeHg concentrations during the growth of the strain ISS312 in experiments at 5 μM were taken at different incubation times. Samples were also collected for the 1 μM growth curves, but as similar degradation kinetics were observed for both concentrations (**Supplementary Table S7**), we have only presented the results from the 5 μM growth curve where we were able to detect a great capacity to remove Hg by strain ISS312 (**Figure 4B**). Hence, in samples taken at 6 h and 12 h, which corresponded to the lag phase of the growth curve, we observed that MeHg concentrations were reduced by 36 % (2.6 μM) and 72 % (1.1 μM), respectively (**Figure 4B**). Furthermore, at time 24 h, when almost all MeHg was removed (with a concentration of only 0.07 μM, and a removal of 98.2 %) the microorganism began the exponential growth phase. After 48 h, MeHg could not be detected (**Figure 4B Supplementary Table S7**). Based on the results obtained, it can be hypothesized that during the lag phase, in the presence of MeHg, the *mer* operon machinery is being activated by the MerR protein involved in the regulation of the operon, and MerA and MerB proteins are subsequently transcribed to catalyze, first, the demethylation of MeHg to Hg^2+^ (MerB), and then its transformation to Hg^0^ (MerA), which is then volatilized. Once the levels of Hg compounds have dropped considerably, and are no longer toxic, cells can grow normally reaching standard growth rates and continue to remove MeHg until very low concentrations of inorganic Hg remained, evidencing the high detoxification capability of strain ISS312, comparable to other Hg resistant bacteria characterized for bioremediation strategies (*80*).

We assumed that most of the MeHg was degraded biotically by the *mer* operon encoded in ISS312, but a fraction could also be abiotically removed. In order to check which proportion of MeHg was either biotically or abiotically removed, we took additional samples at times 0 h and 72 h from the strain culture and from different control treatments (a killed control and medium alone) in the presence of MeHg. As expected, in the culture we did not observe any remaining MeHg at 72 h. However, we detected a certain level of abiotic degradation in the medium and killed controls (**Supplementary Table S8**). We found that MeHg concentration was reduced by 25 % in the absence of bacteria, suggesting that three quarters could be removed biotically while the rest could be degraded by abiotic processes. Still, most part of the MeHg transformation to Hg^2+^ and then to volatile Hg^0^ is caused by ISS312 strain by the operation of *merA* and *merB* genes confirming the idea that biotic MeHg degradation would play a major role in the open ocean, and especially in the bathypelagic as this strain was originally isolated from those deep waters.

### 2.5 Global distribution of ISS312 merAB genes in the bathypelagic ocean

The biogeographic and size fraction distribution of ISS312 genome was assessed in all available bathypelagic metagenomes of the Malaspina Expedition since isolate ISS312 was originally retrieved from bathypelagic waters of the South Atlantic Ocean. We found that this strain affiliating to *Alteromonas mediterranea* species was distributed across all the temperate bathypelagic waters, including the Atlantic, the Pacific and the Indian Oceans (**Figure 5**). Its abundances, according to the data from the fragment recruitment analyses (FRA), varied across ocean basins and we found significative differences between the Pacific and the Brazil basins (P-value= 0.019), and between the Pacific and the Canary basins (P-value: 0.011), suggesting a higher abundance of this bacterium in the Atlantic Ocean. Despite finding these differences between oceans, we did not find significative differences between plankton size fractions, indicating that the isolate could be present in both the free-living (0.2-0.8 μm) and particle-attached (0.8-20 μm) bacterial communities (**Figure 6**). This analysis confirmed, at least for *Alteromonas mediterranea*, the prevalence and wide distribution of this bacterial species carrying Hg resistance genes, which can actively degrade the toxic MeHg, across the global bathypelagic ocean, and its occurrence in both plankton size fractions analyzed. In fact, *Alteromonas* genus has been described to be able to live associated to particulate organic matter (*65*, *81*), which forms in surface layers and sinks into the deep ocean. Besides, *merA* and *merB* genes affiliating to *Alteromonas* have been already detected among the microbial communities associated to sinking particles (*64*). In addition, MeHg production, presenting its maximum concentrations in deeper waters (200-1000 m depth) (*14*, *18*, *19*) usually near the thermocline, is linked to microbial remineralization of particulate organic matter (*18–20*, *23*, *33*). Thereby, this study has uncovered that heterotrophic isolated bacteria from genus *Alteromonas* containing *mer* genes are present in the open ocean from different oceanographic regions, depths and plankton size fractions. These taxa carried in their genomes the key genes for the detoxification of Hg toxic compounds. To the best of our knowledge the detection of *mer* genes from non-contaminated sites and in open ocean waters at a large scale has not been previously reported.

**Figure 5.**
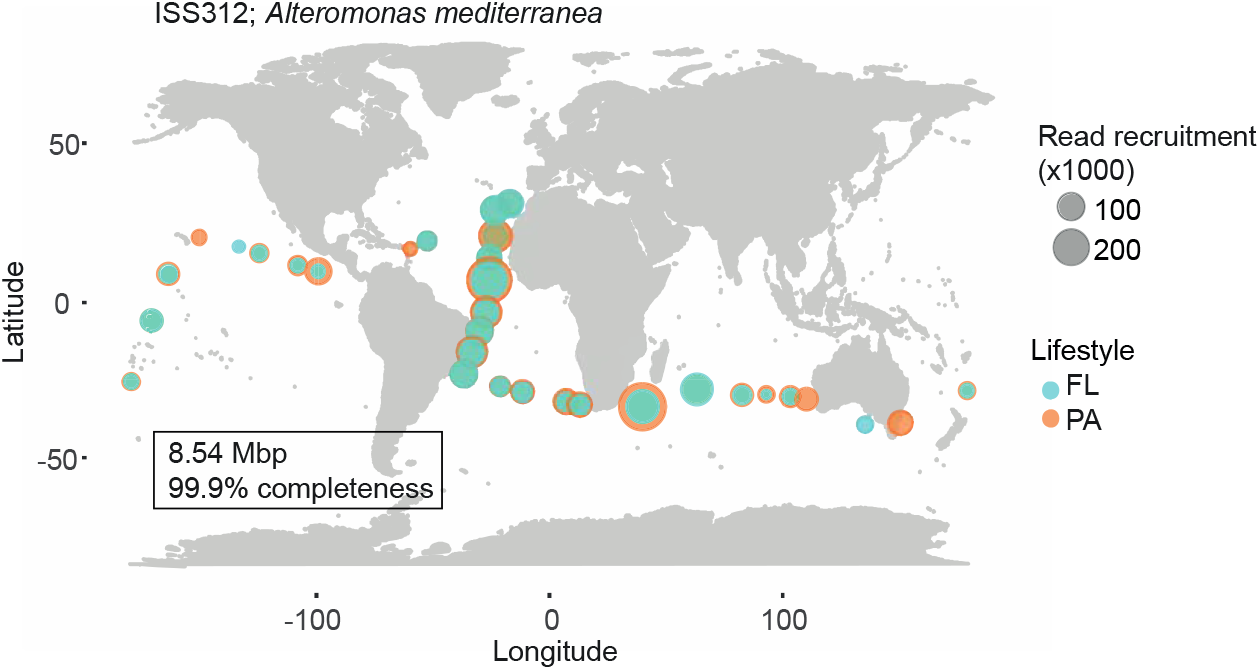
World map showing the distribution of *Alteromonas mediterranea* strain ISS312. Size of the dots indicate number of reads (x1000) and color indicate if the reads were recruited in the free-living (FL, 0.2-0.8 μm) or in the particle-attached (PA, 0.8-20 μm) bacterial communities of the bathypelagic samples.

**Figure 6.**
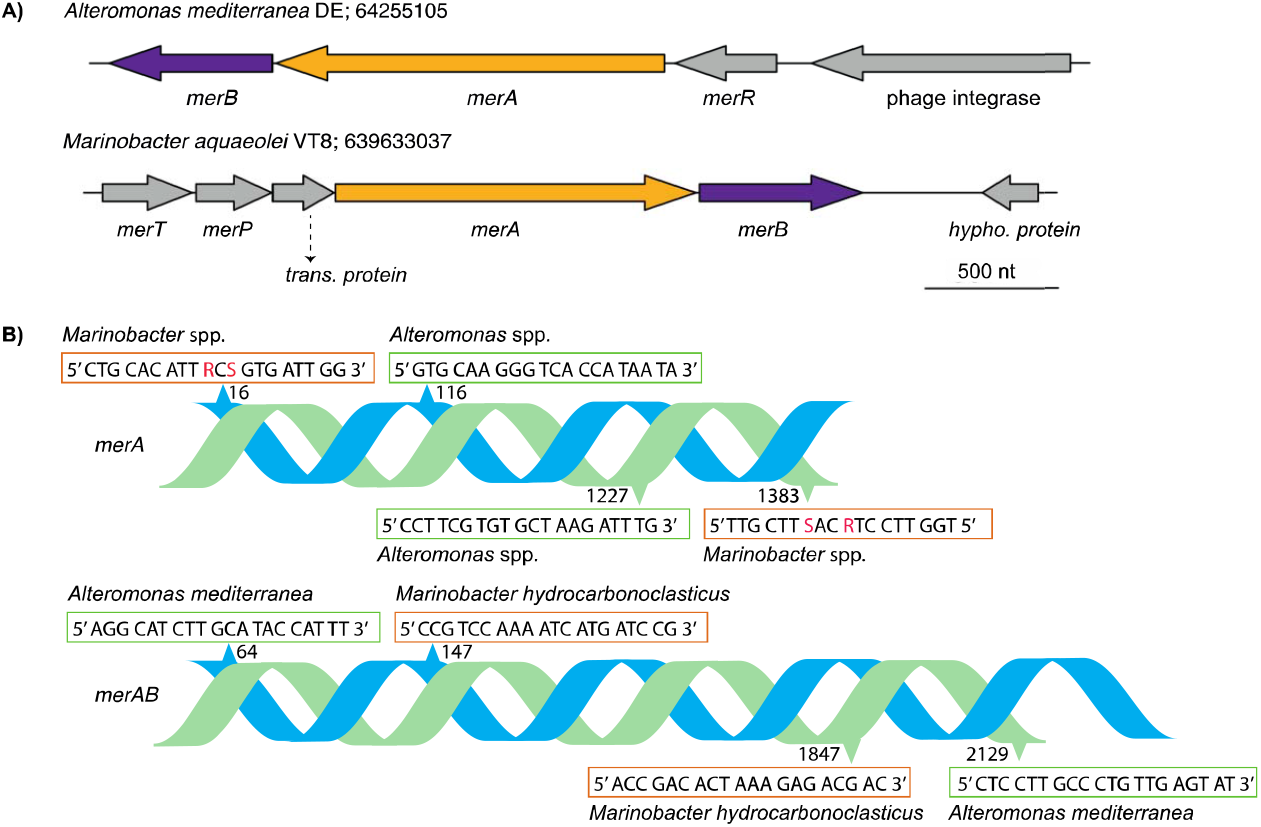
Synteny and primers of *merA* and *merB* genes. **(A)** Co-localization of *merA* and *merB* genes in *Alteromonas mediterranea* DE and *Marinobacter aquaeolei* VT8. Coordinates extracted from the JGI/IMG database. Accession number in the IMG/JGI database is indicated with the species name. Genes: *merA*, mercuric reductase; *merB*, organomercurial lyase; *merR*, *mer* operon regulator; *merT*, *merP* and trans. prot.: mercury transport proteins; hypho. prot: hypothetical protein. **(B)** Forward (blue chain) and reverse (green chain) primers sequences for *merA* and *merAB* genes. Numbers indicate the position where primers attached to the forward and reverse DNA strands. Some of the primers display degenerated bases (see in red): R: A or G; S: G or C.

### 2.6 Outlook

In summary, this study, that nourishes from a large marine bacterial culture collection, offers the first description of the presence and distribution of *merA* and *merB* genes, as well as, the assessment of the tolerance capacity (activity) to mercury compounds, in *Alteromonas* and *Marinobacter* strains, at a broad scale covering different oceanographic regions and depths including the bathypelagic ocean. Hg resistant *Alteromonas* and *Marinobacter* strains were detected in samples from photic and deep ocean waters including the NW Mediterranean Sea, the North and South Atlantic, the Southern and the Indian Oceans. Our results highlighted that mercury resistance genes are widely distributed in the open ocean, and that degradation of the neurotoxic MeHg can be performed through the ocean water column by some specific taxa at high efficiency with important implications in the biogeochemical cycle of mercury and potentially for the environment and human health. Specifically, we have revealed that *Alteromonas mediterranea* strain ISS312 isolated from bathypelagic waters of the South Atlantic Ocean is widely distributed in the global bathypelagic ocean both in the free-living and the particle-attached bacterial communities. Given its high tolerance, the growth rates observed, its efficiency in the removal of MeHg, and its global oceanic distribution, this isolate could be a promising candidate for future Hg marine bioremediation studies.

## 3. Material and Methods

### 3. 1 Selection of marine strains for merAB functional screening

A total of 2003 marine strains were previously isolated from a wide variety of oceanographic regions and depths. Detailed sampling, isolation procedures and partial sequencing of the 16S rRNA gene have been already described in Sanz-Sáez et al. (*35*). This heterotrophic marine bacterial culture collection, called MARINHET, was the basis for the functional screening of Hg resistant bacteria and the description of the distribution of mercury resistance genes of this study. First, we performed a preliminary search of marine genera codifying both *merA* and *merB* in all available finished genomes in the Integrated Microbial Genomes (IMG) database at that time (March 2016) of the Joint Genome Institute (JGI) by: (i) searching the functional annotation of *merA* (as mercuric reductase or mercuric ion reductase) and *merB* (organomercurial lyase or alkylmercury lyase), and (ii) using the Kyoto Encyclopedia of Genes and Genomes (KEGG) orthologs (*82*) K00520 for *merA* and K00221 for *merB*. Secondly, we downloaded the 16S rRNA gene sequences of those genomes in order to BLASTn (*83*) them against the partial 16S rRNA sequences of our isolates and to obtain a list of putative candidates carrying the *merA* and *merB* genes. From those, we focused on two genera, *Alteromonas* and *Marinobacter*, highly abundant taxa in our MARINHET cultured collection. A total 290 isolates from both *Alteromonas* (244) and *Marinobacter* (46) genera were finally selected for functional screening of the Hg resistance genes *merA* and *merB*. These strains originated from a variety of oceanographic regions and depths and covered both the photic and aphotic regions of the water column. Information about the origin of the samples where isolates were retrieved is summarized in **Table 2**.

### 3.2 Primers design for merA and merB genes

Sequences of the *merA* and *merB* genes were downloaded from IMG/JGI. First, we looked for “mercuric reductase” or “mercuric ion reductase” (*merA*) and “organomercurial lyase” or “alkylmercury lyase” (*merB*) genes in the published finished genomes of 22 *Alteromonas* and 6 *Marinobacter* (IMG database in 2016). Then, we downloaded the nucleotide sequences of 15 *merA* and 1 *merB* genes present in those *Alteromonas* genomes, as well as 9 *merA* and 1 *merB* genes from *Marinobacter* (**Supplementary Table S9**). Specific primers pairs were designed separately for: (i) *merA* of *Alteromonas*, (ii) *merA* of *Marinobacter*, (iii) *merA* + *merB* of *Alteromonas*, and (iv) *merA* + *merB* of *Marinobacter*. Both sets of *merA* primers were selected after alignments of the sequences in Clustal Omega (https://www.ebi.ac.uk/Tools/msa/clustalo/) in order to check for the most conserved areas of the gene. For *Alteromonas*, as different *merA* gene copies could be present within the same genome, the designed primers only covered the gene/s copies with the highest similarity between them and between the taxa, since evidence of horizontal gene transfer has been detected in *mer* genes, usually in plasmids, transposons or genomic islands (*65*) **(Supplementary Table S9)**. On the other hand, as *Marinobacter merA* sequences differ greatly from species to species and most of the *Marinobacter* isolates present in our culture collection affiliated to *Marinobacter hydrocarbonoclasticus* or *Marinobacter salarius*, primers were designed to cover the *merA* sequence variants of these species (**Supplementary Table S9**). Otherwise, as *merB* genes were only found in *Alteromonas mediterranea* DE strain (CP003917) and *Marinobacter aquaeolei* VT8 strain (NC_008740) (also named *Marinobacter hydrocarbonoclasticus* VT8), *merA* + *merB* sets of primers were designed with the online tool Primer-BLAST of the National Center for Biotechnology Information (https://www.ncbi.nlm.nih.gov/tools/primer-blast/). The input sequence for the generation of these primers was a concatenate nucleotide sequence of the *merA* and *merB* genes that were co-localized one next to the other in the *mer* operon (**Figure 6A**). All the designed primers had to meet the following requirements: optimum polymerase chain reaction (PCR) product size of 1200 bp for *merA* or 2100 bp for *merA* + *merB* (referred hereafter as *merAB*), annealing temperature around 57 °C, primers length 20 bp, and 50 % GC content. In **Figure 6B** we summarize the sequences of the different sets of primers used in this study.

### 3.3 DNA extraction and PCR conditions

The primers previously designed were used for the screening of *merA* and *merAB* Hg resistance genes in the 290 selected strains. DNA of all the strains was extracted from 48 h liquid cultures grown in Zobell broth medium (i.e. 5 g peptone, 1 g yeast extract in 750 ml of 30 kDa filtered seawater and 250 ml of Milli-Q water) using the DNeasy Blood & Tissue kit (Qiagen) following the manufacturer’s recommendations. Each PCR reaction with a final volume of 25 μl contained: 2 μl of template DNA, 0.5 μl of each deoxynucleotide triphosphate at a concentration of 10 μM, 0.75 μl of MgCl_2_ 1.5 mM, 0.5 μl of each primer at a concentration of 10 μM, 0.125 μl of Taq DNA polymerase (Invitrogen), 2.5 μl of PCR buffer supplied by the manufacturer (Invitrogen, Paisley, UK) and Milli-Q water up to the final volume. Reactions were carried out in a Biorad thermocycler using the following program: initial denaturation at 94 °C for 5 min, followed by 30 cycles of 1 min at 94 °C, 1 min at 55 °C and 2 min at 72 °C, and a final extension step of 10 min at 72 °C. The PCR products were verified and quantified by agarose gel electrophoresis with a standard low DNA mass ladder (Invitrogen). Purification and OneShot Sanger sequencing of *merA* and *merAB* gene products was performed by Genoscreen (Lille, France) with both forward and reverse primers. Geneious software v.11.0.5 (*84*) was used for manual cleaning and quality control of the sequences.

### 3.4 Phylogenetic analyses of the 16S rRNA genes and merA or merAB gene of positive strains

A phylogeny of the isolates screened by PCR for *merA* and *merAB* genes was inferred from their partial 16S rRNA sequences in order to detect a possible clustering between all the positive strains. The closest sequence to each isolate 16S rRNA gene in SILVA v.132 database was found and collected using BLASTn (*83*). Alignment of the isolates and reference sequences was performed with MUSCLE from the Geneious software v.11.0.5 (*84*). The alignment was trimmed to the common 16S rRNA gene fragment covered by both sets of sequences. Phylogeny was constructed using maximum-likelihood inference with RAXML-NG 0.9.0 (*85*) and the GTR evolutionary model with optimization in the among-site rate heterogeneity model and the proportion of invariant sites (GTR+G+I), and 100 bootstrap replicates. In the same way a phylogenetic tree was constructed with the partial 16S rRNA sequences of the positive isolates only. In this tree the closest match in SILVA v.132 database was also included. Presence of *merA* and *merAB* genes, origin of the strains, plus their MIC to HgCl_2_ and MeHg were added with Interactive Tree of Life (ITOL) (*86*).

On the other hand, phylogenetic trees were also constructed with the amino acid sequences of the amplified *merA* and *merAB* genes. As reference sequences we included the best BLASTn hits against UniProtKB sequences retrieved with KEGG identifiers K00520 (*merA*) and K00221(*merB*). Sequences were aligned with ClustalW of the Genious software v.11.0.5 (*84*) with the Gonnet substitution matrix and default gap extension and opening penalties as described previously (*29*). For *merA*, the dihydrolipoamide dehydrogenase protein sequences from *Magnetospirillum magneticum* AMB-1 (WP_011386317.1) and *Pseudomonas fluorescens* Pf0-1 (WP_011336663.1) served as outgroups. Likewise, we included, only in the alignment, the *merA* sequence of *Streptomyces lividans* (P30341), which served to trim the N-terminal region of the aligned sequences such that only the core domain of *merA*, corresponding to positions 1–464 of this *Streptomyces lividans* remained (*87*). In the case of *merAB*, outgroup sequences were not included. Phylogenetic trees were constructed using maximum-likelihood inference with RAXML-NG 0.9.0 (*85*) and the LG evolutionary model with optimization in the among-site rate heterogeneity model and the proportion of invariant sites (LG+G+I), and 100 bootstrap replicates.

### 3.5 Minimum inhibitory concentration experiments

Minimum inhibitory concentration (MIC) assays were designed based on previous studies (*65*, *88*) in order to assess the tolerance of the marine strains to different concentrations of inorganic Hg (as mercury(II) chloride, HgCl_2_) and organic Hg (as methylmercury chloride, CH_3_HgCl) and thus, to test the activity of *merA* and *merB* genes respectively. A stock solution of HgCl_2_ was prepared at 500 μM with autoclaved Milli-Q water. Liquid cultures of the strains growing in Zobell broth with an optical density (O.D. at 600nm) of 0.1 were placed in 24-well plates and inoculated with 5 μM, 10 μM, 20 μM, 25 μM and 50 μM HgCl_2_. In specific cases growth was observed in all HgCl2 concentrations and further MIC assays were done increasing the final concentrations to 50 μM, 60 μM, 70 μM, 80 μM, 90 μM and 100 μM. The tolerance to CH_3_HgCl was also tested for the most tolerant strains to HgCl_2_. In these case, 24-well plates were inoculated with the stock solution to reach final concentrations at 2.5 μM, 5 μM, 10 μM, 15 μM and 20 μM. In all plates a positive control (liquid culture of the strains not amended with CH_3_HgCl or HgCl_2_) and a negative control (liquid culture of a non-resistant marine strain of the genus *Erythrobacter*) were included in the assays. Plates were sealed with parafilm and incubated at room temperature (RT, ~20 °C) and in the dark for 72 h. Visual lectures and O.D. lectures at 600 nm were done in a 24 h period using an automatic plate reader (Infinite ^®^ M200, Tecan) and data was collected using the Magellan^™^ Data Analysis Software (Tecan Diagnostics^®^).

### 3.6 Growth curves

Growth curves were performed to characterize the growth rates of the most tolerant strain (ISS312) to different concentrations of MeHg. We prepared 200 ml of liquid cultures in Zobell broth supplemented with CH_3_HgCl at final concentrations of 0 μM (positive control), 1 μM, 2.5 μM and 5 μM in triplicates. The initial O.D. at 600 nm of the cultures was 0.05 in order to assure enough concentration of cells for growing. Samples for O.D. measurements and for bacterial cell counts were taken approximately every two hours. O.D. was measured at 600 nm with a spectrophotometer (Varian Cary^®^ 100 UV-Vis) and cells were stained with 4’,6-diamidino-2-phenylindole (DAPI) and counted with an automated microscope Zeiss Axio Imager Z2M (*89*, *90*) using the automated image analysis software ACME Tool (www.technobiology.ch). Predicted growth curves based on O.D. observations and kinetics values (growth rates (*μ_max_*), carrying capacity (k) and lag phase time) were calculated with R package *growthcurver v*.0.3.0 (*91*) and GrowthRates v.4.3 software (*92*). For graphical representation, replicates of the different growth curves experiments at several MeHg concentrations were averaged. Hence, mean O.D. and standard deviation was calculated for each time point of the curves.

### 3.7 Measurement of the biotic and abiotic degradation of MeHg

In order to characterize the MeHg degradation rates of our bacteria caused by the action of the *merA* and *merB* genes, 2 ml samples were taken from the 1 μM and 5 μM growth curves of the most tolerant strain, *Alteromonas* sp. ISS312, at times 0, 6, 12, 24 and 48 h. Besides, in order to check the possibility that MeHg was being abiotically removed, we measured the MeHg concentrations from samples taken from multiwell plates experiments. These included one well of a liquid culture from the ISS312 strain in Zobell broth (initial O.D. at 600 nm of 0.05) amended with 5 μM CH^3^HgCl and incubated at RT during 72 h in the dark, as well as two different controls to detect the possible abiotic degradation of MeHg: (i) medium control (CH_3_HgCl at concentrations of 1 μM and 5 μM with no strain added), and (ii) killed control (CH_3_HgCl at concentrations of 1 μM and 5 μM added to autoclaved liquid cultures of the strain). Three replicates of 2 ml samples from each well, containing the CH_3_HgCl treatment and the different controls, were taken at times 0 and 72 h. Collected samples were immediately frozen at -80 °C. Concentration of Hg species was measured by direct derivatization of the culture samples with sodium tetraethyl borate and injection into a hyphenated system consisting of a gas chromatograph coupled to an atomic fluorescence detector via pyrolysis (GC-pyro-AFS) as previously described elsewhere (*93*). Briefly, 2 ml samples were used for derivatization. The pH of the extracts was adjusted to 3.9 by adding 5 ml of 0.1 M acetic acid-sodium acetate buffer and ammonia (20 %) if necessary. Then, 2 ml of hexane and 250 μl of sodium tetraethyl borate (6 %, w/v) were added and the mixture was manually shaken for 5 min. The sample was centrifuged for 5 min at 600 g. The organic layer was transferred to a chromatographic glass vial and stored at -18 °C until analysis. When Hg species were not detectable, the organic layer was preconcentrated under a gentle stream of nitrogen to a low volume (50-100 μL) just before the measurement. The procedural detection limits, after preconcentration, were 0.19 and 0.23 nM for MeHg and inorganic Hg, respectively.

### 3.8 Preparation and observation of samples for transmission electron microscopy

In order to study potential effects of MeHg on strain ISS312 cells morphology, the isolate was grown separately during 24 h in Zobell broth not amended with CH_3_HgCl and amended at a final concentration of 5 μM with shaking in the dark. The overnight culture was centrifuged at 1000 g during 15 min and the supernatant was discarded. The pellet was fixed with paraformaldehyde 2 % final concentration during 30 min at RT. After fixation, the pellet was processed as previously described (*94*) to finally obtain thin sections of the samples that were examined by transmission electron microscopy (TEM, JEM-1400 plus, JEOL). Visualizations were done by the microscopy service of the *Universitat Autònoma de Barcelona* (http://sct.uab.cat/microscopia/en/content/inici).

### 3.9 Strain ISS312 genome sequencing

DNA from strain ISS312 was extracted using the DNeasy Blood & Tissue kit (Qiagen), following the manufacturer’s recommendations. A genome library was prepared with a Celero^™^ DNA-seq library system and sequencing was performed with paired-ended 300 bp long reads by IGA-Tech with a MiSeq Illumina machine. The sequence data was filtered to remove the adapters and the unpaired reads with *cutadapt* v.1.16 (*95*) and the quality was assessed before and after with *fastaqc* v.0.11.7. The clean data were used to do the assembly with Spades v.3.12.0 and then optimized with the same program. QUAST v.5.0.1 (*96*) and ALE were used to assess the quality of the assemblies and the best scores were selected. K-mer 121, 125 and 127 were selected for the optimized-combined assembly and the quality was assessed again. The annotation was done with Prokka v1.13 (*97*) and the completeness of the genome was checked with CheckM v1.0.18 (*98*). In order to search for plasmids within our contigs, we used the database PLSDB (*99*).

### 3.10 Fragment recruitment analysis of the genome of strain ISS312 in bathypelagic metagenomes

The abundance of ISS312 across the global bathypelagic ocean was assessed thanks to its complete genome. Fragment Recruitment Analysis (FRA) was performed by mapping the metagenomic reads of 58 bathypelagic microbial metagenomes from 32 stations (*100*) from Malaspina Expedition, including free-living (0.2-0.8 μm) and particle-attached (0.8-20 μm) bacterial communities. Analysis were done with BLASTn v2.7.1+ (*83*) using the following alignment parameters*: -perc_identity* 70, *-evalue* 0.0001. Only those reads with more than 90 % coverage and mapping identity equal to or higher than 95 % were kept for analysis. In order to remove possible false mapping hits to the conserved regions of rRNA genes, reads aligning to the regions annotated as ribosomal genes were not considered for the analysis. Read counts from mapped reads from each metagenome were corrected by their sequencing depth to make them comparable through samples.

### 3.11 Statistical analyses

ANOVA tests from the *stats* package of the R Statistical Software (*101*) was applied in order to observe possible differences between oceanographic regions in terms of number of positive strains harboring *merA* and/or *merAB* genes. To assess significance, the statistical analyses were set to a conservative alpha value of 0.05. Further, non-parametric Kruskal-Wallis test, from the *stats* package of the R Statistical Software, (*101*) was applied followed by the post hoc pairwise Wilcox test to see the differences between FRA results in different oceanographic regions and between free-living and particle-attached bacterial communities. To assess significance, the statistical analyses were set to a conservative alpha value of 0.05.

### 3.12 Nucleotide accession numbers

Mercury detoxification genes (*merA* and *merAB*) detected in this study through PCR are deposited in GenBank under accession numbers MW273028 - MW273125. *Alteromonas mediterranea* ISS312 genome was deposited in ENA under study accession number PRJEB46669.

## Acknowledgements

We are grateful to Elisabet Laia Sà for helping in the laboratory. We thank the Spanish ministry of Science, Innovation and Universities for granting ISS with a PhD FPU grant (FPU14/03590). We are also grateful to the MER_CLUB project (863584-MER_CLUB-EMFF-BlueEconomy-2018) for hiring ISS in order to finish this study.

## Author contributions

Conceptualization: ISS, SGA, OS

Experimental procedures: ISS, CPG, LT, MPiF, MC, RCRM-D

Data analyses: ISS, PS

Supervision: SGA, OS

Writing: ISS, AGB, SGA, OS

## References

1. M. W. Miller, T. W. Clarkson, Mercury, mercurial, and mercaptans. (Thomas, Springfield, ILL., 1973).

2. T. W. Clarkson, Mercury: major issues in environmental health. Environ. Health Perspect. 100, 31–38 (1993).

3. H. M. Amos, D. J. Jacob, D. G. Streets, E. M. Sunderland, Legacy impacts of alltime anthropogenic emissions on the global mercury cycle. Global Biogeochem. Cycles. 27, 410–421 (2013).

4. UN-Environment, 2019. Global Mercury Assessment 2018. UN-Environment Programme, Chemicals and Health Branch, Geneva, Switzerland. 59 pp.

5. H. H. Eriksen, F. X. Perrez, The Minamata convention: A comprehensive response to a global problem. Rev. Eur. Comp. Int. Environ. Law. 23, 195–210 (2014).

6. A. Saiz-Lopez, S. P. Sitkiewicz, D. Roca-Sanjuán, J. M. Oliva-Enrich, J. Z. Dávalos, R. Notario, M. Jiskra, Y. Xu, F. Wang, C. P. Thackray, E. M. Sunderland, D. J. Jacob, O. Travnikov, C. A. Cuevas, A. U. Acuña, D. Rivero, J. M. C. Plane, D. E. Kinnison, J. E. Sonke, Photoreduction of gaseous oxidized mercury changes global atmospheric mercury speciation, transport and deposition. Nat. Commun. 9, 4796 (2018).

7. M. Enrico, G. Le Roux, N. Marusczak, L.-E. Heimbürger, A. Claustres, X. Fu, R. Sun, J. E. Sonke, Atmospheric mercury transfer to peat bogs dominated by gaseous elemental mercury dry deposition. Environ. Sci. Technol. 50, 2405–2412 (2016).

8. R. P. Mason, G.-R. Sheu, Role of the ocean in the global mercury cycle. Global Biogeochem. Cycles. 16, 1093 (2002).

9. I. Lehnherr, V. L. St. Louis, H. Hintelmann, J. L. Kirk, Methylation of inorganic mercury in polar marine waters. Nat. Geosci. 4, 298–302 (2011).

10. K. M. Munson, C. H. Lamborg, R. M. Boiteau, M. A. Saito, Dynamic mercury methylation and demethylation in oligotrophic marine water. Biogeosciences. 15, 6451–6460 (2018).

11. M. Monperrus, E. Tessier, D. Amouroux, A. Leynaert, P. Huonnic, O. F. X. Donard, Mercury methylation, demethylation and reduction rates in coastal and marine surface waters of the Mediterranean Sea. Mar. Chem. 107, 49–63 (2007).

12. M. Podar, C. C. Gilmour, C. C. Brandt, A. Soren, S. D. Brown, B. R. Crable, A. V. Palumbo, A. C. Somenahally, D. A. Elias, Global prevalence and distribution of genes and microorganisms involved in mercury methylation. Sci. Adv. 1, e1500675 (2015).

13. C. M. Gionfriddo, M. T. Tate, R. R. Wick, M. B. Schultz, A. Zemla, M. P. Thelen, R. Schofield, D. P. Krabbenhoft, K. E. Holt, J. W. Moreau, Microbial mercury methylation in Antarctic sea ice. Nat. Microbiol. 1, 16127 (2016).

14. R. P. Mason, A. L. Choi, W. F. Fitzgerald, C. R. Hammerschmidt, C. H. Lamborg, A. L. Soerensen, E. M. Sunderland, Mercury biogeochemical cycling in the ocean and policy implications. Environ. Res. 119, 101–117 (2012).

15. G. Harding, J. Dalziel, P. Vass, Bioaccumulation of methylmercury within the marine food web of the outer Bay of Fundy, Gulf of Maine. PLoS One. 13, e0197220 (2018).

16. D. Mergler, H. A. Anderson, L. H. M. Chan, K. R. Mahaffey, M. Murray, M. Sakamoto, A. H. Stern, Methylmercury exposure and health effects in humans: A worldwide concern. Ambio. 36, 3–11 (2007).

17. M. R. Karagas, A. L. Choi, E. Oken, M. Horvat, R. Schoeny, E. Kamai, W. Cowell, P. Grandjean, S. Korrick, Evidence on the human health effects of low-level methylmercury exposure. Environ. Health Perspect. 120, 799–806 (2012).

18. D. Cossa, B. Averty, N. Pirrone, The origin of methylmercury in open Mediterranean waters. Limnol. Oceanogr. 54, 837–844 (2009).

19. E. M. Sunderland, D. P. Krabbenhoft, J. W. Moreau, S. A. Strode, W. M. Landing, Mercury sources, distribution, and bioavailability in the North Pacific Ocean: Insights from data and models. Global Biogeochem. Cycles. 23, GB2010 (2009).

20. J. D. Blum, B. N. Popp, J. C. Drazen, C. Anela Choy, M. W. Johnson, Methylmercury production below the mixed layer in the North Pacific Ocean. Nat. Geosci. 6, 879–884 (2013).

21. C. R. Hammerschmidt, K. L. Bowman, Vertical methylmercury distribution in the subtropical North Pacific Ocean. Mar. Chem. 132-133, 77–82 (2012).

22. E. G. Malcolm, J. K. Schaefer, E. B. Ekstrom, C. B. Tuit, A. Jayakumar, H. Park, B. B. Ward, F. M. M. Morel, Mercury methylation in oxygen deficient zones of the oceans: No evidence for the predominance of anaerobes. Mar. Chem. 122, 11–19 (2010).

23. C. H. Lamborg, C. R. Hammerschmidt, K. L. Bowman, An examination of the role of particles in oceanic mercury cycling. Philos. Trans. R. Soc. A. 374, 20150297 (2016).

24. T. Zhang., H. Hsu-Kim, Photolytic degradation of methylmercury enhanced by binding to natural organic ligands Tong. Nat. Geosci. 3, 473–476 (2010).

25. P. Seller, C. A. Kelly, J. W. M. Rudd, A. R. Mac Hutchon, Photodegradation of methylmercury in lakes. Nature. 380, 694–697 (1996).

26. M. Costa, P. S. Liss, Photoreduction of mercury in sea water and its possible implications for Hg0 air-sea fluxes. Mar. Chem. 68, 87–95 (1999).

27. L. J. Stal, M. S. Cretoiu, Chapter 1. What is so special about marine microorganisms? Introduction to the marine microbiome - from diversity to biotechnological applications in The Marine Microbiome (Springer, Switzerland, 2016). pp.3–20.

28. T. Barkay, S. M. Miller, A. O. Summers, Bacterial mercury resistance from atoms to ecosystems. FEMS Microbiol. Rev. 27, 355–384 (2003).

29. E. S. Boyd, T. Barkay, The mercury resistance operon: From an origin in a geothermal environment to an efficient detoxification machine. Front. Microbiol. 3, 349 (2012).

30. R. S. Oremland, C. W. Culbertson, M. R. Winfrey, Methylmercury decomposition in sediments and bacterial cultures: involvement of methanogens and sulfate reducers in oxidative demethylation. Appl. Environ. Microbiol. 57, 130–137 (1991).

31. K. L. Bowman, R. E. Collins, A. M. Agather, C. H. Lamborg, C. R. Hammerschmidt, D. Kaul, C. L. Dupont, G. A. Christensen, D. A. Elias, Distribution of mercury-cycling genes in the Arctic and equatorial Pacific Oceans and their relationship to mercury speciation. Limnol. Oceanogr. 65, S310–S320 (2019).

32. F. Wang, R. W. Macdonald, D. A. Armstrong, G. A. Stern, Total and methylated mercury in the Beaufort Sea: The role of local and recent organic remineralization. Environmetal Sci. Technol. 46, 11821–11828 (2012).

33. L.-E. Heimbürger, D. Cossa, J.-C. Marty, C. Migon, B. Averty, A. Dufour, J. Ras, Methyl mercury distributions in relation to the presence of nano-and picophytoplankton in an oceanic water column (Ligurian Sea, North-western Mediterranean). Geochim. Cosmochim. Acta. 74, 5549–5559 (2010).

34. B. Gworek, O. Bemowska-Kałabun, M. Kijeńska, J. Wrzosek-Jakubowska, Mercury in Marine and Oceanic Waters—a Review. Water, Air, Soil Pollut. 227, 371 (2016).

35. I. Sanz-Sáez, G. Salazar, P. Sánchez, E. Lara, M. Royo-Llonch, E. L. Sà, T. Lucena, M. J. Pujalte, D. Vaqué, C. M. Duarte, J. M. Gasol, C. Pedrós-Alió, O. Sánchez, S. G. Acinas, Diversity and distribution of marine heterotrophic bacteria from a large culture collection. BMC Microbiol. 20, 207 (2020).

36. K. Nakamura, J. Aoki, M. Yamamoto, Mercury volatilization by the most mercury-resistant bacteria from the seawater of Minamata Bay in various physiological conditions. Clean Prod. Process. 2, 174–178 (2000).

37. J. De, N. Ramaiah, Characterization of marine bacteria highly resistant to mercury exhibiting multiple resistances to toxic chemicals. Ecol. Indic. 7, 511–520 (2007).

38. W. Zhang, L. Chen, D. Liu, Characterization of a marine-isolated mercury-resistant Pseudomonas putida strain SP1 and its potential application in marine mercury reduction. Appl. Microbiol. Biotechnol. 93, 1305–1314 (2012).

39. M. Jayaprakashvel, S. Vijay, C. P. Karthigeyan, A. J. Hussain, Isolation and characterization of mercury resistant bacteria from the coastal are of Chennai, India. Int. J. Adv. Res. 4, 64–76 (2015).

40. H.-H. Chiu, W. Y. Shieh, S. Y. Lin, C.-M. Tseng, P.-W. Chiang, I. Wagner-Dö Bler, Alteromonas tagae sp. nov. and Alteromonas simiduii sp. nov., mercury-resistant bacteria isolated from a Taiwanese estuary. Int. J. Syst. Evol. Microbiol. 57, 1209–1216 (2007).

41. X. Deng, P. Wang, Isolation of marine bacteria highly resistant to mercury and their bioaccumulation process. Bioresour. Technol. 121, 342–347 (2012).

42. G. O. Oyetibo, S. T. Ishola, W. Ikeda-Ohtsubo, K. Miyauchi, M. O. Ilori, G. Endo, Mercury bioremoval by Yarrowia strains isolated from sediments of mercury-polluted estuarine water. Appl. Microbiol. Biotechnol. 99, 3651–3657 (2015).

43. C. Vetriani, Y. S. Chew, S. M. Miller, J. Yagi, J. Coombs, R. A. Lutz, T. Barkay, Mercury adaptation among bacteria from a deep-sea hydrothermal vent. Appl. Environ. Microbiol. 71, 220–226 (2005).

44. K. Nakamura, M. Iwahara, K. Furukawa, Screening of organomercurial-volatilizing bacteria in the mercury-polluted sediments and seawater of Minamata Bay in Japan. Clean Prod. Process. 3, 104–107 (2001).

45. A. A. Lima de Silva, M. A. R. de Carvalho, S. A. L. de Souza, P. M. T. Dias, R. G. da Silva Filho, C. S. de Meirelles Saramago, C. A. de Melo Bento, E. Hofer, Heavy metal tolerance (Cr, Ag AND Hg) in bacteria isolated from sewage. Brazilian J. Microbiol. 43, 1620–1631 (2012).

46. B. H. Olson, T. Barkay, R. R. Colwell, Role of plasmids in mercury transformation by bacteria isolated from the aquatic environment. Appl. Environ. Microbiol. 38, 478–485 (1979).

47. T. Barkay, Adaptation of aquatic microbial communities to Hg stress. Appl. Environ. Microbiol. 53, 2725–2732 (1987).

48. J. Simbahan, E. Kurth, J. Schelert, A. Dillman, E. Moriyama, S. Jovanovich, P. Blum, Community analysis of a mercury hot spring supports occurrence of domain-specific forms of mercuric reductase. Appl. Environ. Microbiol. 71, 8836–8845 (2005).

49. L. D. Rasmussen, C. Zawadsky, S. J. Binnerup, G. Oregaard, S. J. Sørensen, N. Kroer, Cultivation of hard-to-culture subsurface mercury-resistant bacteria and discovery of new merA gene sequences. Appl. Environ. Microbiol. 74, 3795–3803 (2008).

50. M. Zeyaullah, B. Islam, A. Ali, Isolation, identification and PCR amplification of merA gene from highly mercury polluted Yamuna river. African J. Biotechnol. 9, 3510–3514 (2009).

51. M. O. Fashola, V. M. Ngole-Jeme, O. O. Babalola, Heavy metal pollution from gold mines: Environmental effects and bacterial strategies for resistance. Int. J. Environ. Res. Public Health. 13, 1047 (2016).

52. A. Ciok, K. Budzik, M. K. Zdanowski, J. Gawor, J. Grzesiak, P. Decewicz, R. Gromadka, D. Bartosik, L. Dziewit, Plasmids of psychrotolerant polaromonas spp. isolated From Arctic and Antarctic glaciers - diversity and role in adaptation to polar environments. Front. Microbiol. 9, 1285 (2018).

53. J. De, N. Ramaiah, A. Mesquita, X. N. Verlekar, Tolerance to various toxicants by marine bacteria highly resistant to mercury. Mar. Biotechnol. 5, 185–193 (2003).

54. A. K. Møller, T. Barkay, M. A. Hansen, A. Norman, L. H. Hansen, S. J. Sørensen, E. S. Boyd, N. Kroer, Mercuric reductase genes (merA) and mercury resistance plasmids in High Arctic snow, freshwater and sea-ice brine. FEMS Microbiol. Ecol. 87, 52–63 (2014).

55. L. Baumann, P. Baumann, M. Mandel, R. D. Allen, Taxonomy of aerobic marine eubacteria. J. Bacteriol. 110, 402–429 (1972).

56. H. Eilers, J. Pernthaler, F. O. Glöckner, R. Amann, Culturability and in situ abundance of pelagic Bacteria from the North Sea. Appl. Environ. Microbiol. 66, 3044–3051 (2000).

57. M. M. Floyd, J. Tang, M. Kane, D. Emerson, Captured diversity in a culture collection: case study of the geographic and habitat distributions of environmental isolates held at the american type culture collection. Appl. Environ. Microbiol. 71, 2813–2823 (2005).

58. A. Gärtner, M. Blümel, J. Wiese, J. F. Imhoff, Isolation and characterisation of bacteria from the Eastern Mediterranean deep sea. Antonie van Leeuwenhoek, Int. J. Gen. Mol. Microbiol. 100, 421–435 (2011).

59. K. M. Handley, J. R. Lloyd, Biogeochemical implications of the ubiquitous colonization of marine habitats and redox gradients by Marinobacter species. Front. Microbiol. 4, 136 (2013).

60. I. Lekunberri, J. M. Gasol, S. G. Acinas, L. Gómez-Consarnau, B. G. Crespo, E. O. Casamayor, R. Massana, C. Pedrós-Alió, J. Pinhassi, The phylogenetic and ecological context of cultured and whole genome-sequenced planktonic bacteria from the coastal NW Mediterranean Sea. Syst. Appl. Microbiol. 37, 216–228 (2014).

61. W. Kai, Y. Peisheng, M. Rui, J. Wenwen, S. Zongze, Diversity of culturable bacteria in deep-sea water from the South Atlantic Ocean. Bioengineered. 8, 572–584 (2017).

62. E. Singer, E. A. Webb, W. C. Nelson, J. F. Heidelberg, N. Ivanova, A. Pati, K. J. Edwards, Genomic potential of Marinobacter aquaeolei, a biogeochemical “opportunitroph.”Appl. Environ. Microbiol. 77, 2763–2771 (2011).

63. M. López-Pérez, A. Gonzaga, A.-B. Martin-Cuadrado, O. Onyshchenko, A. Ghavidel, R. Ghai, F. Rodriguez-Valera, Genomes of surface isolates of Alteromonas macleodii: the life of a widespread marine opportunistic copiotroph. Sci. Rep. 2, 696 (2012).

64. K. M. Fontanez, J. M. Eppley, T. J. Samo, D. M. Karl, E. F. DeLong, Microbial community structure and function on sinking particles in the North Pacific Subtropical Gyre. Front. Microbiol. 6, 469 (2015).

65. E. Ivars-Martinez, A.-B. Martin-Cuadrado, G. D’Auria, A. Mira, S. Ferriera, J. Johnson, R. Friedman, F. Rodriguez-Valera, Comparative genomics of two ecotypes of the marine planktonic copiotroph Alteromonas macleodii suggests alternative lifestyles associated with different kinds of particulate organic matter. ISME J. 2, 1194–1212 (2008).

66. B. V. Mathema, B. C. Thakuri, M. Sillanpää, Bacterial mer operon-mediated detoxification of mercurial compounds: a short review. Arch. Microbiol. 193, 837–844 (2011).

67. J. K. Schaefer, J. Yagi, J. R. Reinfelder, T. Cardona, K. M. Ellickson, S. Tel-Or, T. Barkay, Role of the bacterial organomercury lyase (MerB) in controlling methylmercury accumulation in mercury-contaminated natural waters. Environ. Sci. Technol. 38, 4304–4311 (2004).

68. D. Cossa, L.-E. Heimbürger, D. Lannuzel, S. R. Rintoul, E. C. V. Butler, A. R. Bowie, B. Averty, R. J. Watson, T. Remenyi, Mercury in the Southern Ocean. Geochim. Cosmochim. Acta. 75, 4037–4052 (2011).

69. R. K. Math, H. M. Jin, J. M. Kim, Y. Hahn, W. Park, E. L. Madsen, C. O. Jeon, Comparative genomics reveals adaptation by Alteromonas sp. SN2 to marine tidal-flat conditions: Cold tolerance and aromatic hydrocarbon metabolism. PLoS One. 7, e35784 (2012).

70. H. G. Griffin, T. J. Foster, S. Silver, T. K. Misra, Cloning and DNA sequence of the mercuric- and organomercurial-resistance determinants of plasmid pDU1358. Proc. Natl. Acad. Sci. U. S. A. 84, 3112–3116 (1987).

71. S. Mindlin, G. Kholodii, Z. Gorlenko, S. Minakhina, L. Minakhin, E. Kalyaeva, A. Kopteva, M. Petrova, O. Yurieva, V. Nikiforov, Mercury resistance transposons of Gram-negative environmental bacteria and their classification. Res. Microbiol. 152, 811–822 (2001).

72. A. M. Osborn, K. D. Bruce, P. Strike, D. A. Ritchie, Distribution, diversity and evolution of the bacterial mercury resistance (mer) operon. FEMS Microbiol. Rev. 19, 239–262 (1997).

73. L. Bass, C. A. Liebert, M. D. Lee, A. O. Summers, D. G. White, S. G. Thayer, J. J. Maurer, Incidence and characterization of integrons, genetic elements mediating multiple-drug resistance, in avian Escherichia coli. Antimicrob. Agents Chemother. 43, 2925–2929 (1999).

74. M. Harada, K. Ito, N. Nakajima, S. Yamamura, M. Tomita, H. Suzuki, S. Amachi, Genomic analysis of Pseudomonas sp. strain SCT, an iodate-reducing bacterium isolated from marine sediment, reveals a possible use for bioremediation. G3. 9, 1321–1329 (2019).

75. A. K. Møller, T. Barkay, W. A. Al-Soud, S. J. Sørensen, H. Skov, N. Kroer, Diversity and characterization of mercury-resistant bacteria in snow, freshwater and sea-ice brine from the High Arctic. FEMS Microbiol. Ecol. 75, 390–401 (2011).

76. K. Morishita, K. Nakamura, K. Tuchiya, K. Nishimura, M. Iwahara, O. Yagi, Removal of methylmercury from a fish broth by Alteromonas macledii isolated from Minamata Bay. Japanese J. Water Treat. Biol. 42, 45–51 (2006).

77. C. Lamborg, K. Bowman, C. Hammerschmidt, C. Gilmour, K. Munson, N. Selin, C. M. Tseng, Mercury in the anthropocene ocean. Oceanography. 27, 76–87 (2014).

78. R. Zheng, S. Wu, N. Ma, C. Sun, Genetic and physiological adaptations of marine bacterium Pseudomonas stutzeri 273 to mercury stress. Front. Microbiol. 9, 682 (2018).

79. J. B. Robinson, O. H. Tuovinen, Mechanisms of microbial resistance and detoxification of mercury and organomercury compounds: physiological, biochemical, and genetic analyses. Microbiol. Rev. 48, 95–124 (1984).

80. K. Saranya, A. Sundaramanickam, S. Shekhar, S. Swaminathan, T. Balasubramanian, Bioremediation of mercury by Vibrio fluvialis screened from industrial effluents. Biomed Res. Int. 2017, 6509648 (2017).

81. M. Mestre, C. Ruiz-González, R. Logares, C. M. Duarte, J. M. Gasol, M. M. Sala, Sinking particles promote vertical connectivity in the ocean microbiome. Proc. Natl. Acad. Sci. U. S. A. 115, E6799–E6807 (2018).

82. M. Kanehisa, S. Goto, KEGG: kyoto encyclopedia of genes and genomes. Nucleic Acids Res. 28, 27–30 (2000).

83. S. F. Altschul, W. Gish, W. Miller, E. W. Myers, D. J. Lipman, Basic local alignment search tool. J. Mol. Biol. 215, 403–410 (1990).

84. M. Kearse, R. Moir, A. Wilson, S. Stones-Havas, M. Cheung, S. Sturrock, S. Buxton, S. Cooper, S. Markowitz, C. Duran, T. Thierer, B. Ashton, P. Meinties, A. Drummond, Geneious Basic: an integrated and extendable desktop software platform for the organization and analysis of sequence data. Bioinformatics. 28, 1647–1649 (2012).

85. A. M. Kozlov, D. Darriba, T. Flouri, B. Morel, A. Stamatakis, RAxML-NG: a fast, scalable and user-friendly tool for maximum likelihood phylogenetic inference. Bioinformatics. 35, 4453–4455 (2019).

86. I. Letunic, P. Bork, Interactive Tree Of Life (iTOL) v4: recent updates and new developments. Nucleic Acids Res. 47, W256–W259 (2019).

87. T. Barkay, K. Kritee, E. Boyd, G. Geesey, A thermophilic bacterial origin and subsequent constraints by redox, light and salinity on the evolution of the microbial mercuric reductase. Environ. Microbiol. 12, 2904–2917 (2010).

88. I. Wiegand, K. Hilpert, R. E. W. Hancock, Agar and broth dilution methods to determine the minimal inhibitory concentration (MIC) of antimicrobial substances. Nat. Protoc. 3, 163–175 (2008).

89. M. Zeder, A. Ellrott, R. Amann, Automated sample area definition for high-throughput microscopy. Cytom. Part A. 79A, 306–310 (2011).

90. M. Zeder, J. Pernthaler, Multispot live-image autofocusing for high-throughput microscopy of fluorescently stained bacteria. Cytom. Part A. 75A, 781–788 (2009).

91. K. Sprouffske, A. Wagner, Growthcurver: an R package for obtaining interpretable metrics from microbial growth curves. BMC Bioinformatics. 17, 172 (2016).

92. B. G. Hall, H. Acar, A. Nandipati, M. Barlow, Growth rates made easy. Mol. Biol. Evol, 232–238 (2014).

93. J. J. Berzas Nevado, R. C. Rodríguez Martín-Doimeadios, E. M. Krupp, F. J. Guzmán Bernardo, N. Rodríguez Fariñas, M. Jiménez Moreno, D. Wallace, M. J. Patiño Ropero, Comparison of gas chromatographic hyphenated techniques for mercury speciation analysis. J. Chromatogr. A. 1218, 4545–4551 (2011).

94. C. Lee, J. Y. Kim, W. Il Lee, K. L. Nelson, J. Yoon, D. L. Sedlak, Bactericidal effect of zero-valent iron nanoparticles on Escherichia coli. Environ. Sci. Technol. 42, 4927–4933 (2008).

95. M. Martin, Cutadapt removes adapter sequences from high-throughput sequencing reads. EMBnet.journal. 17, 10–12 (2021).

96. A. Gurevich, V. Saveliev, N. Vyahhi, G. Tesler, QUAST: quality assessment tool for genome assemblies. Bioinformatics. 29, 1072–1075 (2013).

97. T. Seemann, Prokka: rapid prokaryotic genome annotation. Bioinformatics. 30, 2068–2069 (2014).

98. D. H. Parks, M. Imelfort, C. T. Skennerton, P. Hugenholtz, G. W. Tyson, CheckM: assessing the quality of microbial genomes recovered from isolates, single cells, and metagenomes. Genome Res. 25, 1043–1055 (2015).

99. V. Galata, T. Fehlmann, C. Backes, A. Keller, PLSDB: a resource of complete bacterial plasmids. Nucleic Acids Res. 47, D195–D202 (2019).

100. C. M. Duarte, Seafaring in the 21St Century: The Malaspina 2010 Circumnavigation Expedition. Limnol. Oceanogr. Bull. 24, 11–14 (2015).

101. R core team, R core team. A language and environment for statistical computing. R foundation for statistical computing, Vienna, Austria https://www.R-project.org/ (2017).

