## Supplementary Material for "Prevalence of heterotrophic methylmercury detoxifying bacteria across oceanic regions"

**This PDF file includes:**

Supplementary Text

Figs. S1 to S2

Headings Supplementary Tables S1 to S9

References (XX to XX)

**Other Supplementary Material for this manuscript include the following:**

Excel file with Supplementary Tables S1 to S9.

**Supplementary Text**

*1. Ribosomal phylogenetic reconstruction of strains codifying merA and/or merB genes*

The phylogenetic analyses of the 16S rRNA gene from all of the detected strains harboring *merA* and *merAB* genes, together with the rest of the screened strains, revealed some patterns (**Supplementary Figure S2**). First, most of the isolated *Alteromonas* strains with *merA* were related to *Alteromonas australica* and *Alteromonas mediterranea*, and only strains affiliating to the last one presented both genes *merA* and *merB (merAB)*. We also detected one strain with *merAB* genes affiliated to *Alteromonas macleodii.* Secondly, *Marinobacter* strains displaying *merAB* were related to *Marinobacter hydrocarbonoclasticus, Marinobacter salarius* and some uncultured *Marinobacter* strains. We are aware that our primers do not match the whole diversity within *Alteromonas* since there is a large uncultured *Alteromonas* cluster related to NW Mediterranean, Indian Ocean and North and South Atlantic samples and from photic and aphotic layers that was not covered by our primer-set (**Supplementary** **Figure S2**). Moreover, this phylogeny showed that positive strains for *merA* and *merB* genes clustered together with strains which do not harbor these genes (**Supplementary** **Figure S2**). It has been described that some bacterial species codify different sequence variants of the *merA* gene (*29*, *74*) including the *Alteromonas* genus (*65*). Therefore, it is possible that some of the *Alteromonas* and *Marinobacter* strains within the MARINHET collection tested presented different sequence variants, although we were only able to detect the ones targeted by the primers designed. On the other hand, the operon *mer* can be either codified in the chromosome (*69*) or in plasmids (*70*, *28*), and usually, *mer* genes are components of transposons (*71*), and integrons (*72*, *73*). Thus, it is not surprising to find some strains within the same species without the *mer* operon.

**Supplementary Figures**

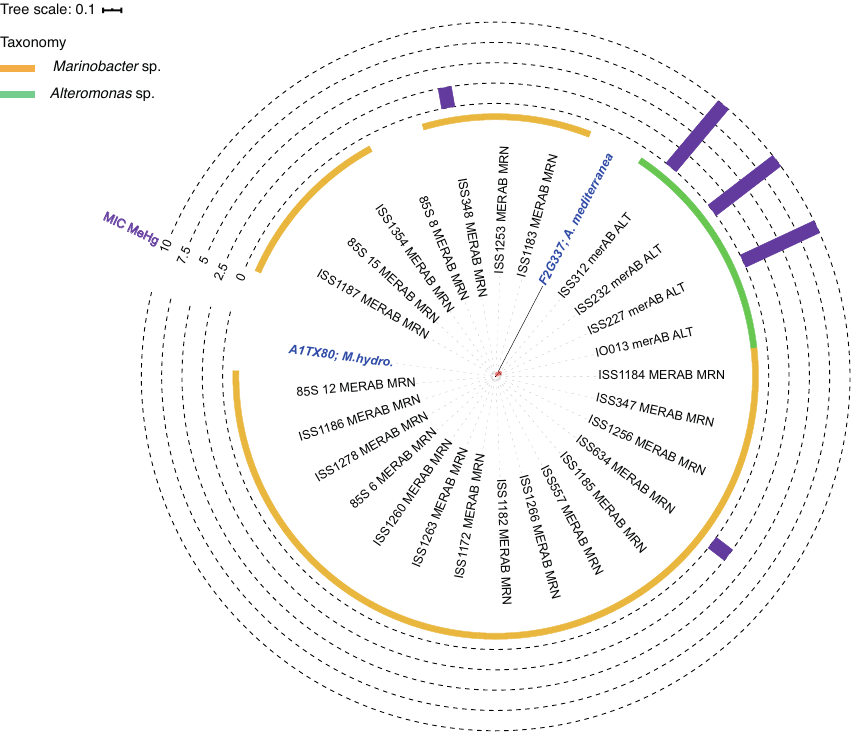

**Figure S1. Phylogenetic tree inferred with the *merAB* amino acid sequences.** Color strip indicates taxonomy of the sequences**:** *Alteromonas* sp., green; *Marinobacter* sp., orange. Reference sequences are indicated in bold blue: *A. mediterranea, Alteromonas mediterranea; M. hydro., Marinobacter hydrocarbonoclasticus.* MIC results for the tested strains against MeHg are indicated by bars. Bootstrap values >75% are indicate by red circles in the tree nodes.

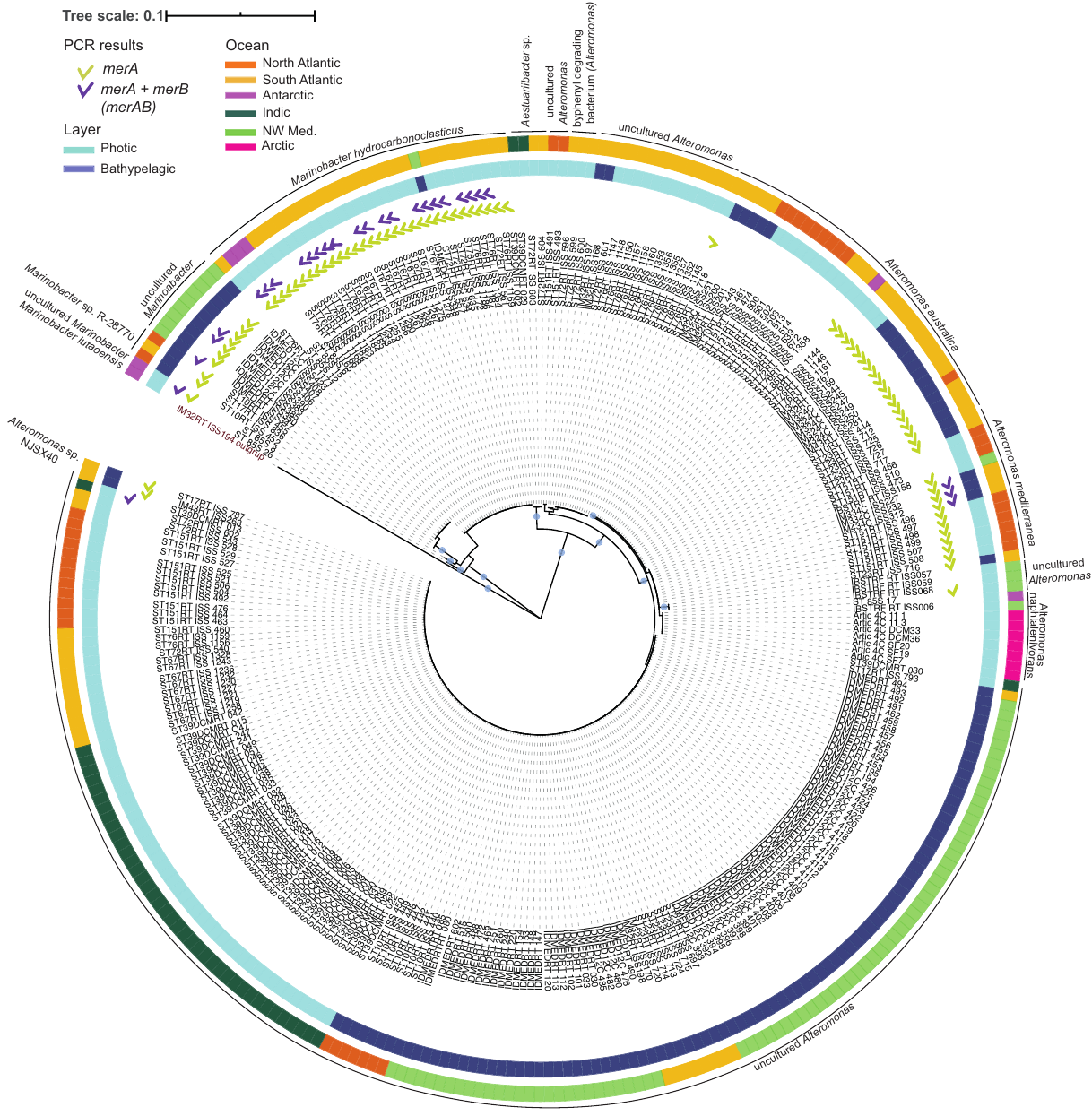

**Figure S2. Phylogenetic tree with the 16S rRNA gene sequences of all the isolates PCR screened for *merA* and *merAB* genes.** Presence or absence of both *merA* and *merAB* genes is indicated by tick symbols. Depth (first color strip) and oceanographic location (second color strip) where the isolates were retrieved are indicated by colors. Bootstrap values ≥ 75 are indicated by red circles in the tree nodes. For graphic representation reference 16S rRNA sequences were removed from original tree, and names of these references sequences had been placed surrounding tree labels.

**Headings Supplementary Tables**

**Table S1.** Candidate genera for mercury bioremediation obtained from the search of *merA* and *merB* genes in the KEGG database and from the IMG/JGI database.

**Table S2.** Summary of the putative isolates within the MARINHET culture collection that could harbor *merA* and *merB* genes based on the BLASTn search between the partial 16S rRNA genes sequences of the isolates and the 16S rRNA gene sequences of the candidate genera extracted from the JGI/IMG database.

**Table S3.** Information of the *Alteromonas sp.* and *Marinobacter* sp. strains used for PCR screening of the *merA* and *merB* genes.

**Table S4.** Summary of the number of positive strains per oceanographic region.

**Table S5.** OTUs defined at 99% clustering with the *Alteromonas* and *Marinobacter* positive strains in order to select candidates for MIC analyses.

**Table S6.** Summary of the results for the MIC determination assays to HgCl_2_.

**Table S7.** Concentrations of MeHg and inorganic mercury at different time points during the growth curves at 1 µM and 5 µM. SD: standard deviation; LOD: limit of detection.

**Table S8.** Biotic (culture) and abiotic controls (killed and medium alone) of the 5 µM MeHg degradation performed by the ISS312 *Alteromonas* strain. LOD: limit of detection.

**Table S9.** Downloaded *merA* and *merB* sequences from the JGI/IMG database (2016) to design the PCR primers. In green are selected those *merA* and *merB* copies that should be recognized by the primers designed.
